# EBF1 regulates the blood-thymus barrier and ETP influx through Claudin-5 in thymic endothelial cells

**DOI:** 10.64898/2026.01.22.700802

**Authors:** Aurelie Lenaerts, Rudolf Grosschedl

## Abstract

The blood-thymus-barrier (BTB) is essential for regulating the entry of thymic-seeding progenitors into the thymus, an organ lacking self-renewing T cell progenitors. Here, we identify Early B-cell factor 1 (*Ebf1*), a transcription factor with established roles in hematopoiesis, as specifically expressed in thymic endothelial cells. Tie2-Cre-mediated deletion of *Ebf1* results in a significant reduction of early thymic progenitors (ETPs), despite unaltered bone marrow lymphoid progenitor populations, unaltered thymic stromal subset frequencies, and intact downstream thymocyte differentiation. Transcriptome analysis of *Ebf1*-deficient thymic endothelial cells reveals increased expression of tight-junction-associated genes, in particular *Cldn5*, a key tight-junction component that restricts the BTB. Consistent with reduced BTB permeability, ETP subset analysis demonstrates a proportional reduction in the ETP1 fraction, the progenitor population that resides in closest proximity to thymic endothelial cells at the cortico-medullary junction. Together, our study implicates *Ebf1* in the regulation of the BTB permeability, through modulation of tight-junction programs, thereby enabling thymus-seeding progenitor entry.

## INTRODUCTION

T cell differentiation occurs in the thymus and is a highly ordered program [1]. Successful T cell receptor (TCR) rearrangement allows for the differentiation of CD4^-^CD8^-^ double-negative (DN) T cells into CD4^+^CD8^+^ double-positive (DP) T cells, followed by positive and negative selection mediated by thymic epithelial cells (TECs) and thymic dendritic cells. These processes result in the generation of mature, single-positive (SP) CD4 and CD8 cells [2]. The thymus lacks self-renewing T cell progenitors, and is dependent on the homing of thymic-seeding progenitors (TSPs) from the bone marrow [3,4]. TSPs can be derived from the continuum of lymphoid progenitors in the BM [5–11], which is represented by MPP4/LMPPs and common lymphoid progenitors (CLPs).

Although research into thymic stromal cell function has mainly been focused on TEC function, non-epithelial thymic stromal cells also play important roles in T cell homeostasis [12,13]. In particular, thymic endothelial cells (ECs) are instrumental for the homing of thymic seeding progenitors (TSP) to the thymus, by expressing specific adhesion molecules and chemokines. TSPs first undergo tethering and rolling on endothelial cells using P-selectin/PSGL-1 interactions, followed by CCL25-induced CCR9 signaling. This results in the rapid activation of αLβ2 and α4β1 integrins, which upon binding with ICAM-1 and VCAM-1, respectively, mediates firm arrest of rolling cells and entry into the thymus [14,15]. In addition, thymic ECs are suggested to act as gatekeepers for thymic homing, by adjusting its CCL25/P-selectin expression, depending on TSP numbers and plasma sphingosine-1-phosphate levels [16]. TSPs that enter the thymus at the cortico-medullary-junction (CMJ) are called early thymic progenitors (ETPs), and are distinguished from other thymic progenitors by cKit expression.

Early B cell Factor 1 (EBF1) is a transcription factor that is expressed in a number of tissues, such as B lymphoid cells [17,18], adipose tissue [19,20], neurons [21] and endothelial cells [22]. Its function has mainly been investigated in the context of transcriptional regulation of B cell differentiation, where it plays a lineage-instructive role [17,23]. *Ebf1* expression in bone marrow mesenchymal stromal cells was also found to regulate cell adhesion and chemotaxis programs, affecting HSC quiescence and self-renewal capacity [22]. Analogous to the bone marrow niche affecting hematopoiesis, in this study we explore how *Ebf1* deletion in the thymus microenvironment affects T lymphopoiesis. We characterized T cell development in the bone marrow, the thymus and in the spleen, upon *Tie2*-Cre mediated *Ebf1* deletion, and observed a specific reduction in ETPs upon *Ebf1* deletion. Furthermore, we analyzed the thymic stromal compartment and *Ebf1*-related transcriptome changes in thymic ECs. Finally, we demonstrate increased expression of tight junction regulator associated with blood-thymus-barrier permeability, *Cldn5*, in *Ebf1* deficient thymic ECs. Our findings suggest that EBF1 participates in the regulation of blood-thymus-barrier permeability.

## RESULTS

### Thymic endothelial cells express *Ebf1*

To examine whether *Ebf1* could play a role in thymus microenvironment, we first assessed *Ebf1* expression in publicly available bulk RNA-sequencing datasets of the thymic endothelial cells (ECs), medullary-and cortical-thymic epithelial cells (mTEC and cTECs) [24]. We find that *Ebf1* is specifically expressed in thymic ECs (**Fig. 1A**). To validate the expression pattern of *Ebf1* in the thymus stromal compartment, we performed quantitative reverse transcriptase (qRT)-PCR for *Ebf1* on sorted TEC and ECs. Consistent with the bulk RNA-sequencing results, qRT-PCR analysis of *Ebf1* expression showed strong expression in ECs and no expression in TECs (**Fig. 1B**). Together, these data suggest that *Ebf1* is specifically expressed in thymic ECs.

**Figure 1.**
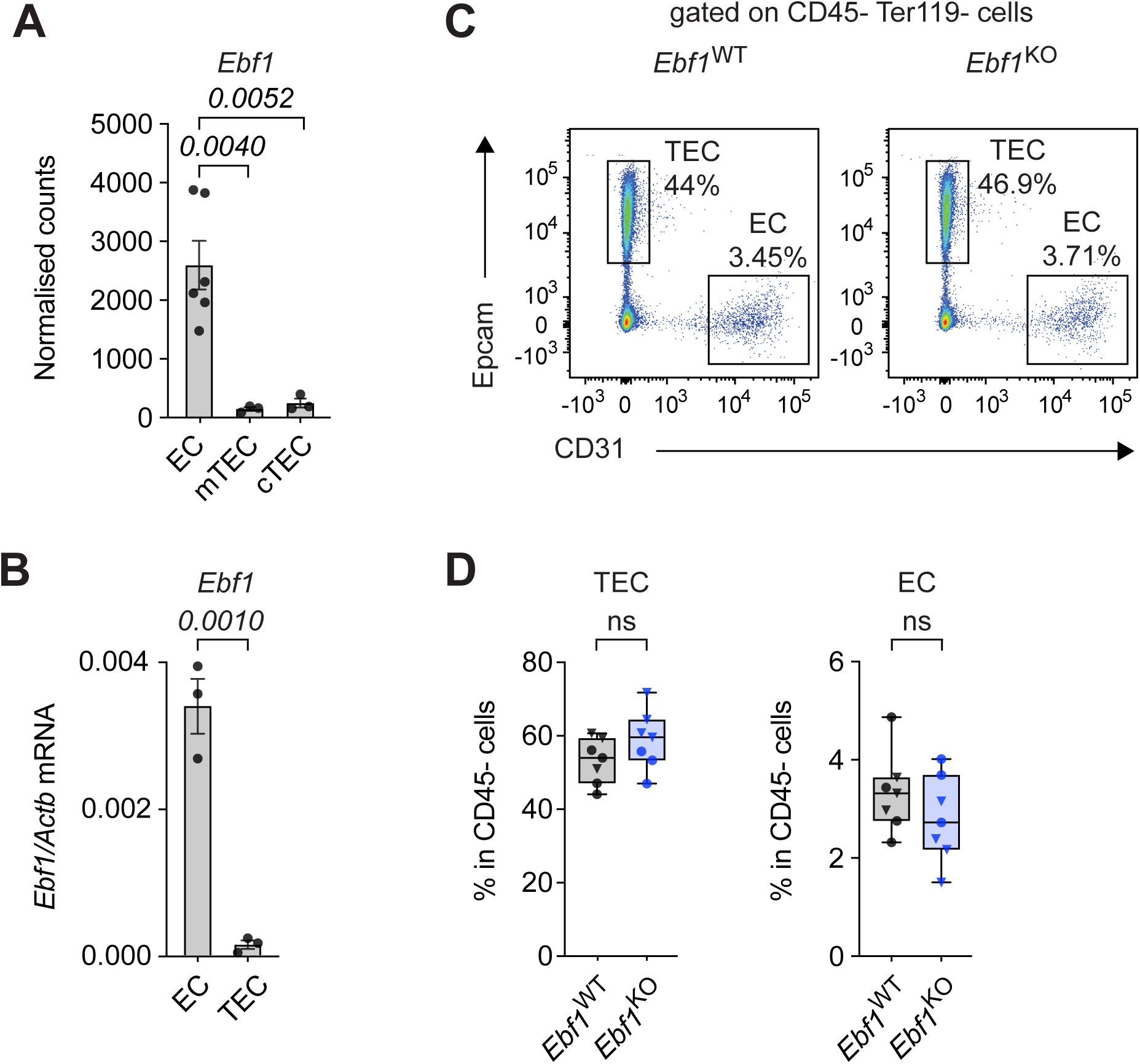
Thymic endothelial cells express *Ebf1*. **(A)** Normalized read counts of *Ebf1* in different thymic stromal populations from public RNA-sequencing dataset [original data from [24]]. In this study a distinction was made between KitL^+^ and KitL^-^ thymic EC cells which have been represented a single endothelial cell (EC) population in this graph. Statistical significance was determined by one-way ANOVA. **(B)** Quantitative reverse-transcriptase (qRT) PCR of *Ebf1* relative to *Actb* mRNA expression. Data are represented as mean ± sem. n=3. Statistical significance was determined by unpaired t-test. **(C)** Representative flow cytometry plots of TECs (CD45^-^Ter119^-^Epcam^+^) and ECs (CD45^-^Ter119^-^CD31^+^) in *Ebf1*^WT^ and *Ebf1*^KO^ mice. **(D)** Frequency of TECs and thymic ECs in CD45-thymic stromal cells. *Ebf1*^WT^ n=7, *Ebf1*^KO^ n=7. Mice were analyzed at 4-6-weeks-old, (•) represent female mice, (▾) represent male mice. Data are represented as boxplots. Statistical significance was determined by unpaired t-test. Data are from ≥ 2 independent experiments. mTEC, medullary thymic epithelial cells; cTEC, cortical thymic epithelial cells; ECs, endothelial cells.

To examine the role of *Ebf1* in thymic ECs on T lymphopoiesis, we conditionally deleted *Ebf1* in endothelial cells, using *Ebf1*^fl/fl^ *Tie2^+/Cre^* mice, hereafter called *Ebf1*^KO^ mice. As a control we used *Ebf1*^wt/wt^ *Tie2^+/Cre^* mice, hereafter called *Ebf1*^WT^ mice. *Tie2^+/Cre^* drives Cre expression in the endothelium and in adult HSCs [25–27]. To examine whether *Ebf1* deletion affects the thymic stromal compartment, we assessed the frequency of thymic ECs and TECs within the CD45^-^ thymic stromal compartment. We observed no significant differences in the frequency of both populations (**Fig. 1C and 1D**), indicating that Tie2-Cre mediated *Ebf1* deletion does not affect the proportion of thymic ECs in the thymic microenvironment.

### *Ebf1* deficiency results in decreased early thymic progenitors in the thymus

To explore the relationship between thymic ECs and T lymphopoiesis upon *Ebf1* deletion, we first assessed early T cell development in the thymus of 4-6-week-old young *Ebf1*^WT^ and *Ebf1*^KO^ mice. We observed no difference in thymus cellularity upon *Ebf1* deletion (**Fig. S1A**). No significant changes were observed in the number of mature single positive (SP) CD4 or CD8 cells, in the number of TCR^+^ double positive (DP) cells, nor in the number of double negative (DN) cells (**Fig. S1C and S1D**). Similarly, in 12-week-old adult mice we observed no differences in thymic cellularity, SP, DP or DN thymic cells between *Ebf1*^WT^ and *Ebf1*^KO^ mice (**Fig. S1B and S1E**). However, we detected a significant decrease in the frequency and absolute number of early thymic progenitors (ETPs; CD4^-^CD8^-^Lin^-^cKit^+^) in the thymus of young (**Fig. 2A and 2B**) and adult (**Fig. S1F and 1G**) *Ebf1*^KO^ mice compared to *Ebf1*^WT^ mice. The DN thymocyte population can be further separated by CD44 and CD25 surface markers into DN1-4 progenitor populations (**Fig. 2C**). Given that ETPs are the seeding progenitors of the thymus, we next examined the subsequent T cell differentiation intermediates. Although a significant decrease was observed in ETP population, remarkably this decrease in progenitors was normalized by the DN1 stage, and no significant changes were observed in the absolute numbers of DN1, DN2, DN3 or DN4 populations in young and adult *Ebf1*^KO^ mice relative to *Ebf1*^WT^ mice (**Fig. 2D and Fig. S1H**). Together, these data indicate that T cell differentiation in the thymus carries on unaffected in *Ebf1*^KO^ mice, even though the seeding ETP population is significantly reduced.

**Figure 2.**
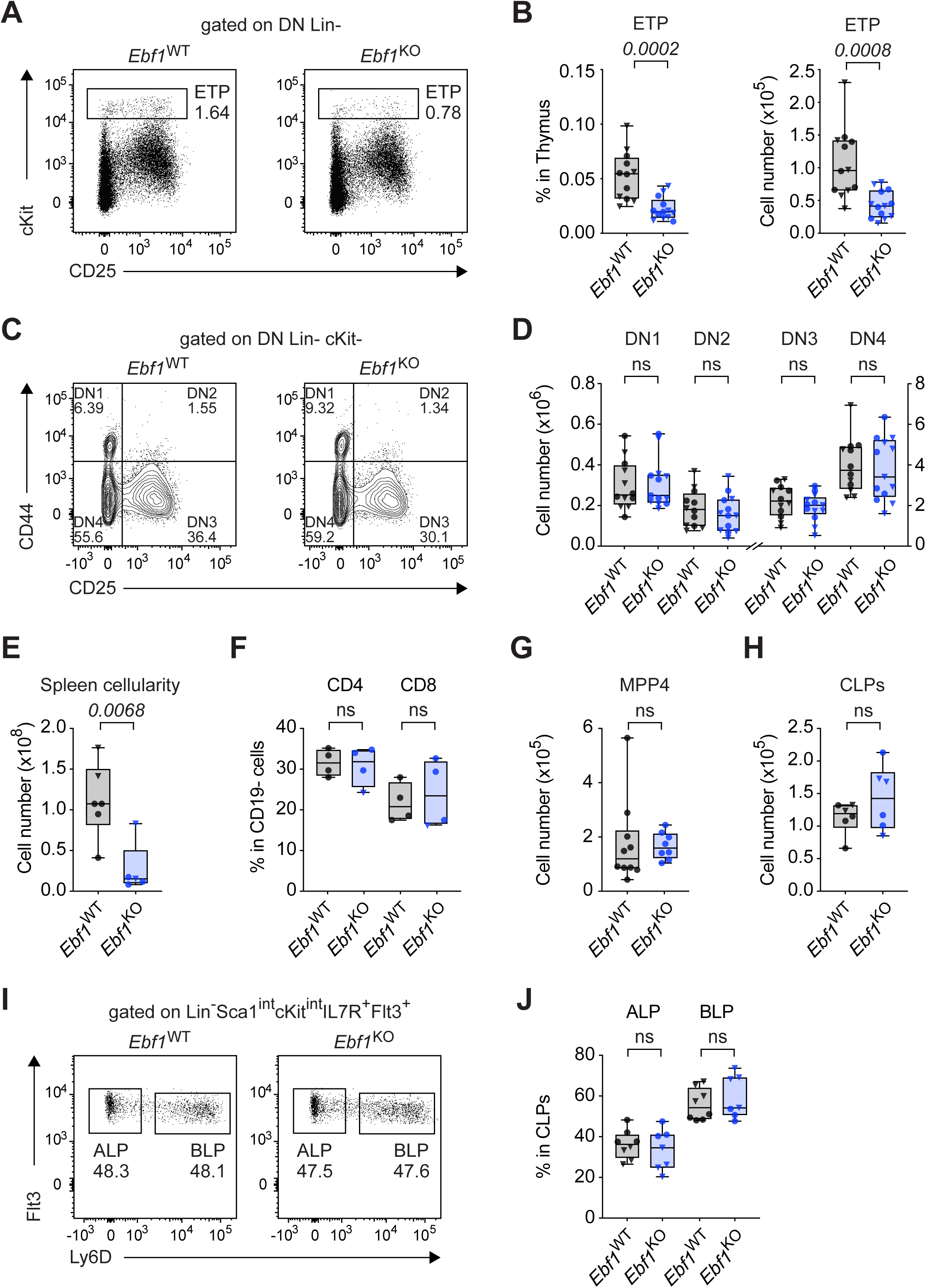
Reduced ETPs upon *Ebf1* deletion. **(A)** Representative dot plots of ETPs (CD4^-^CD8^-^Lin^-^cKit^+^) in *Ebf1*^WT^ and *Ebf1*^KO^ mice. **(B)** Total frequency (left) and absolute number (right) of ETPs in the thymus. **(C)** Representative contour plots for the different DN populations in *Ebf1*^WT^ and *Ebf1*^KO^ mice. **(D)** Absolute number of DN1-4 populations in the thymus. **(B, D)** *Ebf1*^WT^ n=12, *Ebf1*^KO^ n=13. **(E)** Total spleen cellularity. *Ebf1*^WT^ n=6, *Ebf1*^KO^ n=5. **(F)** Frequency of CD4 and CD8 T cells within CD19^-^ cells in the spleen. *Ebf1*^WT^, *Ebf1*^KO^ n=4. **(G)** Absolute number of lymphoid-biased MPP4 (Lin^-^Sca1^+^cKit^+^CD150^-^CD48^+^Flt3^+^) progenitors in the bone marrow. *Ebf1*^WT^ n=10, *Ebf1*^KO^ n=8. **(H)** Absolute number of lymphoid progenitors CLP (Lin^-^Sca1^int^cKit^int^IL7R^+^Flt3^+^) progenitors in the bone marrow. *Ebf1*^WT^, *Ebf1*^KO^ n=6. **(I)** Representative dot plots of ALP and BLP populations in *Ebf1*^WT^ and *Ebf1*^KO^ mice. **(J)** Proportion of ALP and BLP cells in the CLP population. *Ebf1*^WT^ n=8, *Ebf1*^KO^ n=7. (A-J) Mice were analyzed at 4-6-weeks-old, (•) represent female mice, (▼) represent male mice. Data are represented as boxplots. Statistical significance was determined by unpaired t-test. Data are from ≥ 2 independent experiments. ETPs, early thymic progenitors; DN, double negative (CD4^-^CD8^-^Lin^-^cKit^-^).

### Normal T cell maturation in the spleen of *Ebf1*^KO^ mice

In addition to regulating thymic seeding progenitors (TSP) entry, thymic ECs also play a key role in regulating the egress of SP thymocytes into the peripheral lymphoid organs, through the regulation of intrathymic sphingosine-1-phosphate (S1P) levels [28,29]. To examine whether endothelial deletion of *Ebf1* affects thymic egress, we assessed the T cell compartment in the spleen. Consistent with the loss of B cells in *Ebf1*^KO^ mice due to the Tie2-Cre mediated hematopoietic deletion of *Ebf1*, we observed a significant decrease in total spleen cellularity in *Ebf1*^KO^ compared to *Ebf1*^WT^ mice (**Fig. 2E**). In order to analyze the T cell compartment in a meaningful way, we disregarded the loss of B cells and compared CD4^+^ and CD8^+^ T cells within the CD19-negative spleen compartment of *Ebf1*^WT^ and *Ebf1*^KO^ mice. We observed no significant changes in the frequency of CD4 or CD8 T cells in the spleen of *Ebf1*^WT^ and *Ebf1*^KO^ mice (**Fig. 2F**). Together these data suggest that endothelial *Ebf1* deletion does not affect the egress of thymocytes and the maturation of T cells in the periphery.

### Bone marrow lymphoid progenitors are unaffected

Self-renewing lymphoid progenitors are found in the bone marrow (BM), and exist along a continuum from which TSPs can arise. Therefore, we examined the different BM lymphoid progenitors upstream of the ETP population. We observed a similar total BM cellularity in *Ebf1*^KO^ and *Ebf1*^WT^ mice (**Fig. S1I**). Moreover, we detected no significant differences in the numbers of multipotent lymphoid progenitor, MPP4, and in the numbers of common lymphoid progenitor (CLPs) in the BM of *Ebf1*^KO^ mice compared to *Ebf1*^WT^ mice (**Fig. 2G and 2H**). CLPs contain all-lymphoid progenitors (ALP; Lin^-^Sca1^int^cKit^int^IL7R^+^Flt3^+^Ly6D^-^) that give rise to B, T and NK cells, as well as B-lymphoid progenitors (BLP; Lin^-^Sca1^int^cKit^int^IL7R^+^Flt3^+^Ly6D^+^) that generate B cells [30,31]. Therefore, we also examined their relative frequencies, and observed no significant changes in ALPs and BLPs frequencies in the BM of *Ebf1*^KO^ compared to *Ebf1*^WT^ mice (**Fig. 2I and 2J**). Together these data indicate that Tie2-Cre mediated *Ebf1* deletion does not affect lymphoid progenitors in the BM, and likely does not account for the reduction of ETPs in the thymus of *Ebf1*^KO^ mice.

### Specific loss of thymic endothelial cell-associated ETP1 cells

Considering that the BM hematopoietic lymphoid progenitors are comparable between *Ebf1*^WT^ and *Ebf1*^KO^ mice, this led us to examine the ETP population in more detail. *Gata3* is the only member of the GATA transcription factor family that is expressed in T cells [32], and is vital for T lymphopoiesis in the thymus [33–37]. *Gata3* is important for ETP generation, T cell commitment and β-selection in DN2-DN4 thymocytes [38]. Therefore, we assessed the GATA3 expression pattern by intracellular flow cytometric analysis of DN1-DN4 populations in *Ebf1*^WT^ and *Ebf1*^KO^ mice (**Fig. 3A and 3B**). In line with previous studies [1,39], we observed a sharp increase in GATA3 expression in DN2 cells (**Fig. 3A and 3B**). No significant changes were detected in the mean fluorescence intensity (MFI) of GATA3 in the DN1-DN4 populations of *Ebf1*^KO^ mice relative to *Ebf1*^WT^ mice (**Fig. 3B**), suggesting that T cell commitment is not affected by the endothelial deletion of *Ebf1*. Moreover, we did not detect changes of GATA3 expression in ETPs between *Ebf1*^WT^ and *Ebf1*^KO^ mice, indicating that the reduction in ETP population in *Ebf1*^KO^ mice is not due to an intrinsic defect in GATA3 expression (**Fig. 3C**).

**Figure 3.**
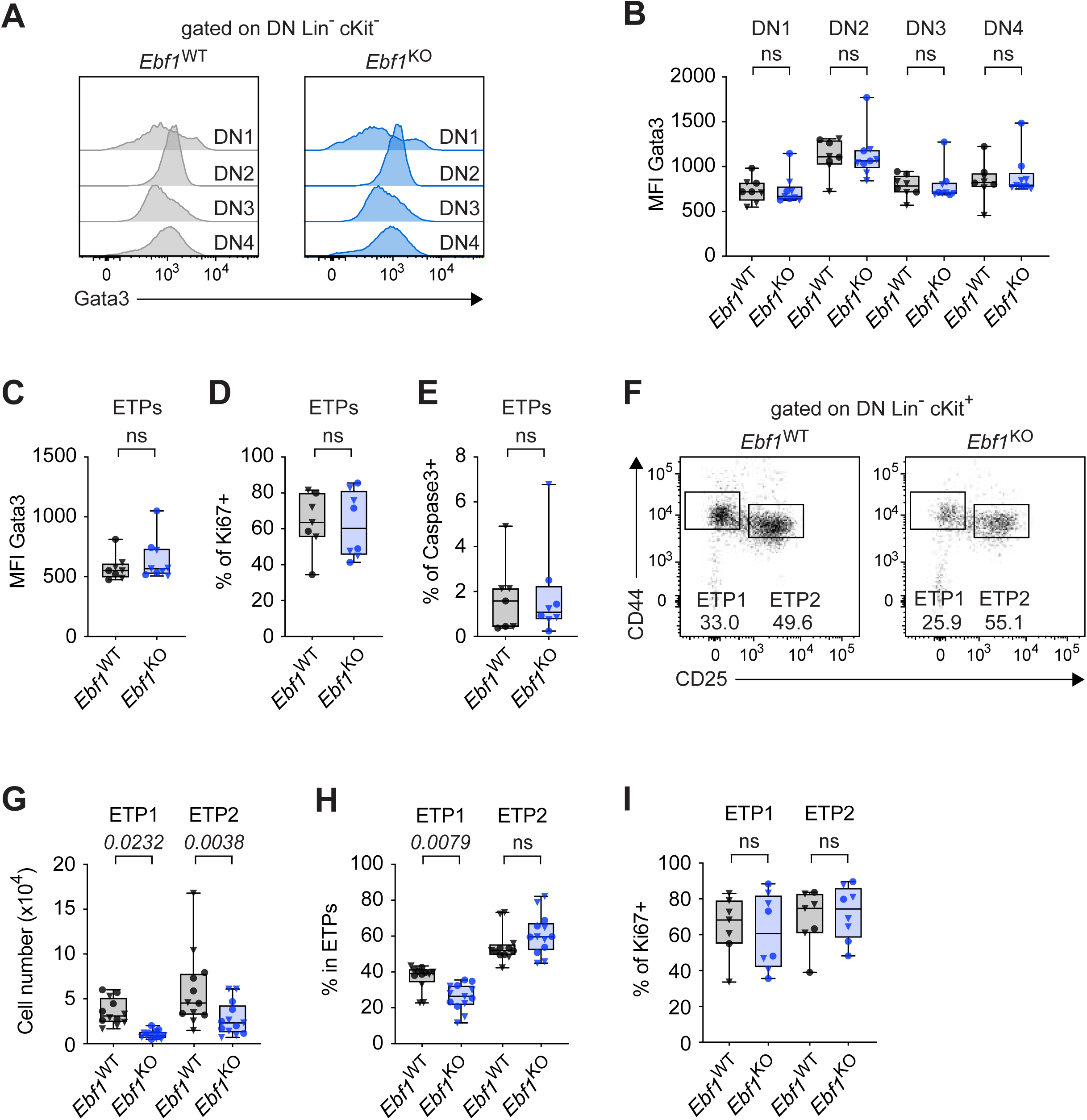
Reduced ETP1 fraction in ETP population upon *Ebf1* deletion. **(A)** Representative histograms of GATA3 expression in DN1-4 populations of the thymus, in *Ebf1*^WT^ and *Ebf1*^KO^ mice. **(B, C)** Mean fluorescence intensity (MFI) of Gata3 **(B)** in DN1-4 populations, and **(C)** in ETPs, in the thymus. *Ebf1*^WT^ n=8, *Ebf1*^KO^ n=9**. (D)** Frequency of Ki67+expressing cells within ETPs. **(E)** Frequency of Caspase3+ expressing cells within ETPs. **(D, E)** *Ebf1*^WT^ n=7, *Ebf1*^KO^ n=8. **(F)** Representative dot plots for ETP1 and ETP2 populations in *Ebf1*^WT^ and *Ebf1*^KO^ mice. **(G)** Absolute number of ETP1 and ETP2 populations in the thymus. **(H)** Frequency of ETP1 and ETP2 cells in the ETP population. **(G, H)** *Ebf1*^WT^ n=12, *Ebf1*^KO^ n=13. **(I)** Frequency of Ki67+ expressing cells within ETP1 and ETP2 cells. *Ebf1*^WT^ n=7, *Ebf1*^KO^ n=8. Mice were analyzed at 4–6-weeks-old, (•) represent female mice, (▼) represent male mice. Data are represented as boxplots. Statistical significance was determined by unpaired t-test. Data are from ≥ 2 independent experiments.

We next considered whether the reduction of ETP cells in *Ebf1*^KO^ mice could be attributed to other intrinsic features, such as decreased proliferation or increased apoptosis. However, we observed no significant differences in the frequency of Ki67 expressing ETPs (**Fig. 3D**), nor in the frequency of Caspase3 expressing ETPs (**Fig. 3E**), in *Ebf1*^KO^ mice compared to *Ebf1*^WT^ mice. The ETP population can be further separated by CD25 expression, into ETP1 and ETP2 populations (**Fig. 3F**). ETP1 cells are similar to DN1 cells in immunophenotype, reside in close proximity to thymic ECs at the cortico-medullary-junction (CMJ), and depend on endothelial cell-derived kitL signal for survival [24,40]. ETP2 cells are similar to DN2 cells in immunophenotype, and are located further into the thymus cortex, away from the thymic ECs at the CMJ. ETP2 cells express cKit and receive kitL signaling from cTECs [24,40]. In line with the significant decrease in ETP numbers in *Ebf1*^KO^ mice relative to *Ebf1*^WT^ mice (**Fig. 2B**), we detected a significant decrease in the numbers of ETP1 and ETP2 cells in *Ebf1*^KO^ mice relative to *Ebf1*^WT^ mice (**Fig. 3G**). Notably, we observed a significant decrease in the proportion of ETP1 cells, and a slight, non-significant increase in the proportion of ETP2 cells within the ETP population, in *Ebf1*^KO^ mice compared to *Ebf1*^WT^ mice (**Fig. 3H**). Though the proliferation status of ETP1 and ETP2 cells are comparable between *Ebf1*^WT^ and *Ebf1*^KO^ mice (**Fig. 3I**). Together these data indicate that *Ebf1* deletion in thymic ECs alters the proportion of ETP1 and ETP2 cells within the ETP population, which was not attributable to intrinsic characteristics such as proliferation status or GATA3 expression levels. These data show a specific decrease in the ETP1 population, which reside in close proximity to thymic ECs, suggesting an impaired interaction between thymic ECs and ETP1.

### Thymic ECs upregulate *Cldn5* upon *Ebf1* deletion

To further understand the molecular effects of *Ebf1* deletion in thymic ECs, we analyzed the transcriptome of thymic ECs from *Ebf1^WT^*and *Ebf1^KO^* mice by bulk RNA-sequencing. We identified 176 differentially regulated genes (FDR < 0.05), of which the majority were downregulated genes (n=131) in *Ebf1*^KO^ relative to *Ebf1*^WT^ thymic ECs (**Fig. 4A**). GO analysis of differentially regulated genes revealed an enrichment of the GO terms, response to TNF, fat cell differentiation and regulation of fibroblast proliferation amongst the genes downregulated in *Ebf1*^KO^ thymic ECs (**Fig. 4B**). Interestingly, we observed a repeated enrichment for vasculature related GO terms amongst the genes upregulated in *Ebf1*^KO^ thymic ECs (**Fig. 4B**). Closer examination of the genes linked to the vasculature signature revealed a downregulation of *Ccl2*, *Gata2* and *Notch4* in *Ebf1*^KO^ thymic ECs, which are critical for endothelial cell function [41,42]. At the same time, we observed an upregulation of genes associated with gap junctions or vascular permeability, such as *Gja5*, *Ddah1* and *Cldn5* (**Fig. 4C**) [43–46].

**Figure 4.**
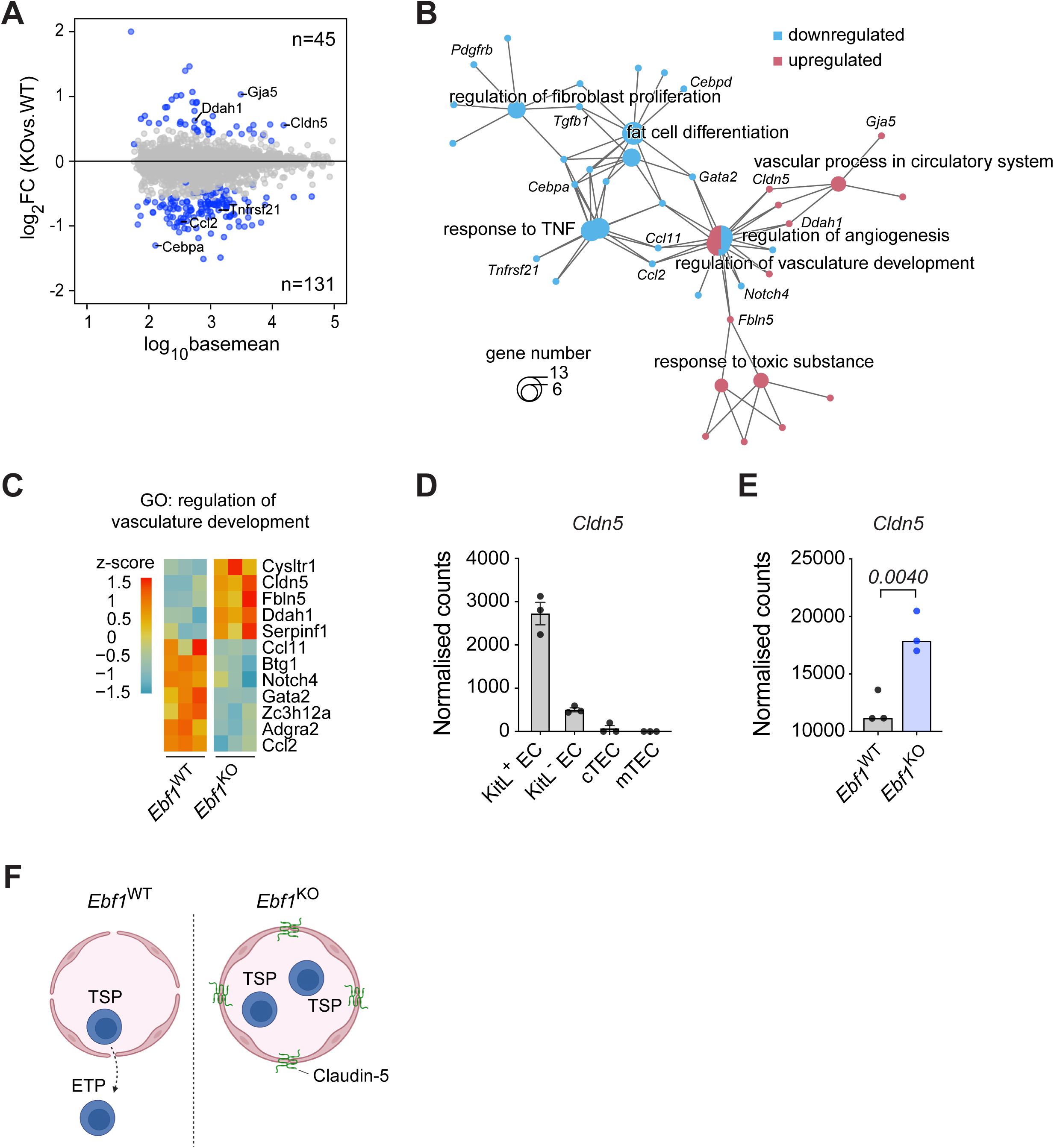
Thymic endothelial cells upregulate *Cldn5* upon *Ebf1* deletion. **(A)** MA plot of thymic ECs comparing *Ebf1*^WT^ and *Ebf1*^KO^ transcriptomes. Dots in blue represent genes with an FDR <0.05. The y axis represents shrunken log_2_ fold change (FC), capped at |log_2_ FC| 2, and the x axis represents log10 basemean. The number of up-and down-regulated genes in *Ebf1*^KO^ mice compared to *Ebf1*^WT^ mice are displayed. **(B)** Cnetplot analysis of DE genes between *Ebf1*^WT^ and *Ebf1*^KO^ thymic ECs. **(C)** Heatmap showing gene expression of DE genes associated with gene-ontology (GO) signature: regulation of vasculature development. Heatmap scale represents z-scores. **(D)** Normalized read counts of *Cldn5* in different thymic stromal populations from public RNA-sequencing dataset [original data from [24]]. **(E)** Normalized read counts of *Cldn5* in *Ebf1*^WT^ and *Ebf1*^KO^ thymic ECs. Mice were analyzed at 4–6-weeks-old, n=3. **(F)** Schematic diagram illustrating reduced permeability of the blood-thymus barrier and impaired trafficking of thymic-seeding progenitors (TSP), associated with increased Claudin-5 expression in *Ebf1^KO^* thymic ECs. EC, endothelial cells; cortical thymic epithelial cells (cTEC); medullary thymic epithelial cells (mTEC); DE, differentially expressed; FDR, false discovery rate.

Gap junctions and tight junctions play a role in regulating blood-tissue barriers, and endothelial tight junctions are constituted of membrane proteins, such as claudins (*Cldn*) [47]. Claudin-5 (*Cldn5*), a member of the claudin membrane protein family, is mainly studied in the context of the blood-brain-barrier and is essential for its establishment [45]. The blood-thymus-barrier (BTB) is found to exist only in the thymic cortex [48,49], whereas the thymic medulla is open to blood circulation [50–52]. A recent study has found that *Cldn5* is preferentially expressed in thymic ECs of the cortex, and proposed that *Cldn5* expression accounts for the differential vascular permeability between the cortex and the medulla [46]. To examine *Cldn5* expression and its relationship with TSP homing, we revisited the public dataset which described KitL^+^ thymic ECs to be the niche for ETPs, and KitL^-^ thymic ECs to be responsible for thymocyte egress [24]. As expected, we find *Cldn5* selectively and highly expressed in KitL^+^ VECs compared to KitL^-^ VECs and TECs (**Fig. 4D**). Modulation of *Cldn5* expression affects the permeability of blood-tissue barriers, and a previous study has shown that overexpression of *Cldn5* in thymic ECs resulted in decreased BTB permeability [46]. In line with this, we observed a significant upregulation of *Cldn5* expression in thymic ECs upon *Ebf1* deletion (**Fig. 4E**). Altogether, our RNA-seq analysis suggests that the permeability of the BTB in *Ebf1*^KO^ mice may be decreased through the upregulation of tight junction genes such as *Cldn5*, thereby resulting in reduced ETP influx into the thymus (**Fig. 4F**).

## DISCUSSION

The role of the thymic microenvironment is mainly investigated in the context of thymic epithelial cells (TECs). In this study, we explored the function of EBF1 in thymic endothelial cells (ECs), and its influence on T lymphopoiesis. We showed that *Ebf1* is specifically expressed in thymic ECs, and that endothelial deletion of *Ebf1* results in a significant decrease of early thymic progenitors (ETPs), with a specific decrease in the thymic EC interacting ETP1 fraction. In addition, we find that *Ebf1*^KO^ thymic ECs downregulated *Gata2* and *Notch4*, which are key regulators for endothelial cell function [41,42]. Finally, we demonstrate increased expression of tight junction regulator, *Cldn5*, in *Ebf1*^KO^ thymic ECs. Increased *Cldn5* expression has been associated with reduced blood-thymus-barrier permeability [46], offering explanation for the decreased ETP population but subsequent normal T lymphopoiesis in *Ebf1*^KO^ mice.

In this study, we make use of the *Tie2*-Cre model to mediate *Ebf1* deletion, thus both hematopoietic and endothelial function must be considered when interpreting results. We observed a pronounced reduction in the ETP population of *Ebf1*^KO^ mice, while subsequent T cell development in the thymus, T cell maturation in the spleen and lymphoid progenitor development in the bone marrow were normal. The normalization of thymic T cell development is not explained by a significant increase in ETP proliferation, nor by a decrease in ETP cell death in *Ebf1*^KO^ compared to *Ebf1*^WT^ mice. It is possible that the retention time of thymocytes in the thymus is longer in *Ebf1*^KO^ compared to *Ebf1*^WT^ mice, however, this is an aspect not further explored in this study. Together, these data indicate that there are no intrinsic defects in T cell development upon *Tie2*-Cre mediated *Ebf1* deletion, in line with the fact that EBF1 expression is absent in T cell lineages, indicating that the reduction in the number of ETPs upon *Ebf1* deletion is mediated via thymic ECs.

While we find that the proportion of ECs in the thymus is unchanged upon *Ebf1* deletion, FACS analysis does not provide information about their distribution and morphology in the thymus. The properties of cortical thymic ECs were shown to be different from medullary thymic ECs, for instance to regulate the blood-thymus-barrier [24,46,49,53]. Furthermore, progenitors that enter the thymus are found in the thymus cortex, implicating cortical ECs in regulating TSP homing. Insight into the molecular consequences of *Ebf1* deletion in thymic ECs was provided by bulk-RNAseq analysis, which showed decreased expression of *Notch4*, and a number of transcription factors, such as C/EBPs (*Cebpa, Cebpb and Cebpe*), and *Gata2*, which play important roles in EC function [41,42,54,55]. This suggests the possibility of additional functional changes in *Ebf1*-deficient thymic ECs, an aspect not further investigated in this study. Notably, we also observed decreased expression of chemokine *Ccl2* and *Cxcl1*, in *Ebf1*^KO^ thymic ECs, which has been shown to increase vascular permeability [56–58]. Furthermore, we observed a significant increase in expression of gap junction factor, *Gja5*, and tight junction member, *Cldn5*, in *Ebf1*^KO^ thymic ECs. *Cldn5* expression is relatively specific to ECs, and is particularly important among the junctional proteins based on the phenotype of *Cldn5*-KO mice, which exhibit extensive vascular permeability of the blood-brain-barrier, leading to lethality soon after birth [45]. Additionally, overexpression of *Cldn5* in an endothelial cell line led to increased transepithelial electrical resistance, and tracer experiments in *Cldn5*-KO mice showed diffuse distribution of the tracer in the thymus cortex compared to WT mice, demonstrating the crucial role of *Cldn5* in establishing the blood-thymus-barrier in the cortex [46]. This led us to hypothesize that increased *Cldn5* expression in *Ebf1*^KO^ mice leads to decreased permeability of the blood-thymus-barrier, and subsequent reduction in the number of ETPs in the thymus (**Fig. 4F**). The regulatory mechanism through which *Ebf1* influences *Cldn5* expression remains unresolved. Notably, *Ebf1* is not differentially expressed between cortical KitL^+^ thymic ECs, which are responsible for TSP homing, and medullary KitL^-^ thymic ECs, which are responsible for thymocyte egress [24].

In conclusion, our findings showed that EBF1 participates in the regulation of BTB permeability in thymic ECs through *Cldn5*, and is of potential interest for the regeneration of the thymus after injury.

## METHODS

### Mice

*Ebf1-flox* mice were generated as detailed in [59]. *Tie2-Cre* [26] mice were purchased from The Jackson Laboratory; stock no. 008863. Only male heterozygous *Tie2^Cre^Ebf1^wt/fl^* mice were used for breeding, as litters with female *Tie2-Cre* drivers demonstrated germline *Ebf1* deletion. flox/flox or +/Cre littermates were used as controls for the *Tie2^Cre^Ebf1^flox^* strain. C57BL/6J mice were bred in-house. 4–14-week-old animals of both sexes were used for experiments. All mice were maintained, bred and analyzed on the C57BL/6J background, in the animal facility of the Max Planck Institute of Immunobiology and Epigenetics under specific pathogen-free conditions. Animals were housed on a 14-hour/10-hour light-dark cycle and provided with standard rodent chow and water ad libitum. All animal procedures were performed in compliance with the approved by responsible Animal Welfare Committees (Regierungspräsidium Freiburg, Karlsruhe/Germany, Nr. 35-9185.81/G-24/116).

### Cell isolation and flow cytometry

To assess the thymus stromal cell populations, the thymus was digested rather than mashed through a cell strainer. Connective tissue was removed from the dissected thymus and minced finely with scissors. The resulting tissue fragments were resuspended in 1 mL RPMI-2%FCS and the supernatant containing thymocytes was discarded. The tissue fragments were then incubated in 1 mL digestion media (RPMI-2%FCS, 0.2mg/ml Col VI, 0.2mg/ml Dispase, 10ug/ml DNaseI) at 37°C for 1h, at 550 rpm. After 30 minutes, the supernatant was collected and added to 5 mL cold RPMI-2%FCS. After the second incubation, 500 μl RPMI-EDTA was added to stop the digestion. To enrich for stromal cell populations, hematopoietic cell depletion was carried out with CD45 Magnetic-activated cell sorting (MACS) beads according to manufacturer’s instructions. CD45-thymic stromal cells were collected and used for downstream applications.

Single cell suspensions were prepared from BM (femora, tibiae and ilia bones) by crushing them in cold PBS-3%FCS using a pestle and mortar. Single cell suspensions were prepared from the thymus and the spleen, by mashing them in cold PBS-3%FCS though a 70µm strainer. Spleen and BM suspensions were subjected to erythrocyte lysis using RBC lysis buffer (BioLegend).

Single cell suspensions were incubated for 30 min with the appropriate antibody cocktail. Intracellular staining was performed by fixing and permeabilizing with the eBioscience Foxp3/Transcription Factor Staining Buffer Set (eBioscience). Surface antibodies were diluted in PBS-3%FCS, intracellular antibodies were diluted in permeabilisation buffer (eBioscience). Between each step, cells were washed twice with PBS-3%FCS or permeabilisation buffer and centrifuged at 400 xg at 4°C. Flow cytometry analysis was performed on an LSRFortessa instrument and cell sorting on FACSAria III instrument (BD Biosciences). Data were analyzed with FlowJo Software V.10 (TreeStar).

### RNA extraction and RT-qPCR

Total RNA was prepared using the Arcturus PicoPure RNA isolation kit (Applied Biosystems) according to manufacturer’s instructions and treated with DNase (Qiagen). cDNA was synthesized with Oligo(dT)_18_ primers using the SuperScript II Reverse Transcriptase Kit (Invitrogen). Quantitative RT-PCR was performed with Taqman Gene Expression Assays (FAM; Thermo Fisher Scientific), with Taqman Fast Advanced Master Mix (Thermo Fisher Scientific) using target probes against *Ebf1* (Mm01288947_g1) normalized to *Act2b* (Mm00607939_s1 Actb). Reactions were performed in duplicate and run on a StepOne Real-Time PCR system (Applied Biosystems).

### Bulk RNA-sequencing

Thymic endothelial cells were sorted into 50 μl of extraction buffer. Total RNA was prepared using the Arcturus PicoPure RNA isolation kit (Applied Biosystems) according to manufacturer’s instructions and treated with DNase (Qiagen). Three biological replicates were used for RNA-seq of thymic endothelial cells from *Ebf1*^WT^ and *Ebf1*^KO^ mice. RNA-seq libraries were prepared by the deep-sequencing facility (MPI-Freiburg) using NEBNext SingleCell/Low Input RNA library prep kit (NEB). Bulk-RNA-seq paired-end 100 bp reads were generated using the Illumina NovaSeq 6000 system, at a depth of 50 million reads.

### Analysis

Raw FASTQ files were mapped against the mm10 reference genome using the mRNA-seq module implemented in snakePipes (version 2.5.1) [60], using the --trim flag. Briefly, reads were aligned using STAR (version 2.7.3a) [61], followed by expression count quantification with featureCounts (version 2.0.0, [62]). Data quality was assessed using Deeptools QC (version 3.3.2, [63]). Genes with an average expression higher than 100 counts in any condition were selected for further analysis. Downstream differential expression analysis was performed using DESeq2 (version 1.28.1) [64] implemented in R (version 4.0.0) and ashr was used for LogFoldChange (LFC) shrinkage of results (version 2.2.47) [65]. Genes were considered differentially expressed at a false discovery rate (FDR) < 0.05. Thymic ECs from *Ebf1*^WT^ and *Ebf1*^KO^ mice were analyzed together. Publicly available data collected for *Ebf1* expression analysis in thymic endothelial cells, were analyzed together [24]. Normalized counts were extracted from the appropriate DEseq2 objects. All figures were generated using the ggplot2 package (version 3.3.0) (Wickham, 2016). Enrichment analysis was performed using clusterProfiler (version 4.10.0) [66].

### Statistical analysis

Data are presented as boxplots or medians, indicated in the figure legends. Details of statistical tests and the exact replicate numbers are reported in the figure legends and/or figures. Except for sequencing analysis, all statistical analyses were performed using Prism 8 software (GraphPad).

### Online supplemental material

**Fig. S1** shows data related to **Fig. 2**.

## AUTHOR CONTRIBUTIONS

A.L designed and performed experiments, bioinformatics analysis. Conceptualization and Writing: A.L, and R.G. Supervision: R.G.

## ACKNOWLEDGMENTS

The authors thank H-J. Schwarz for technical assistance. We thank the MPI-IE core facilities for Deep Sequencing, Mouse and Flow cytometry (especially K. Schuldes), for their invaluable assistance. Work in the Grosschedl Laboratory is supported by funds from the Max Planck Society.

---

**Figure S1.**
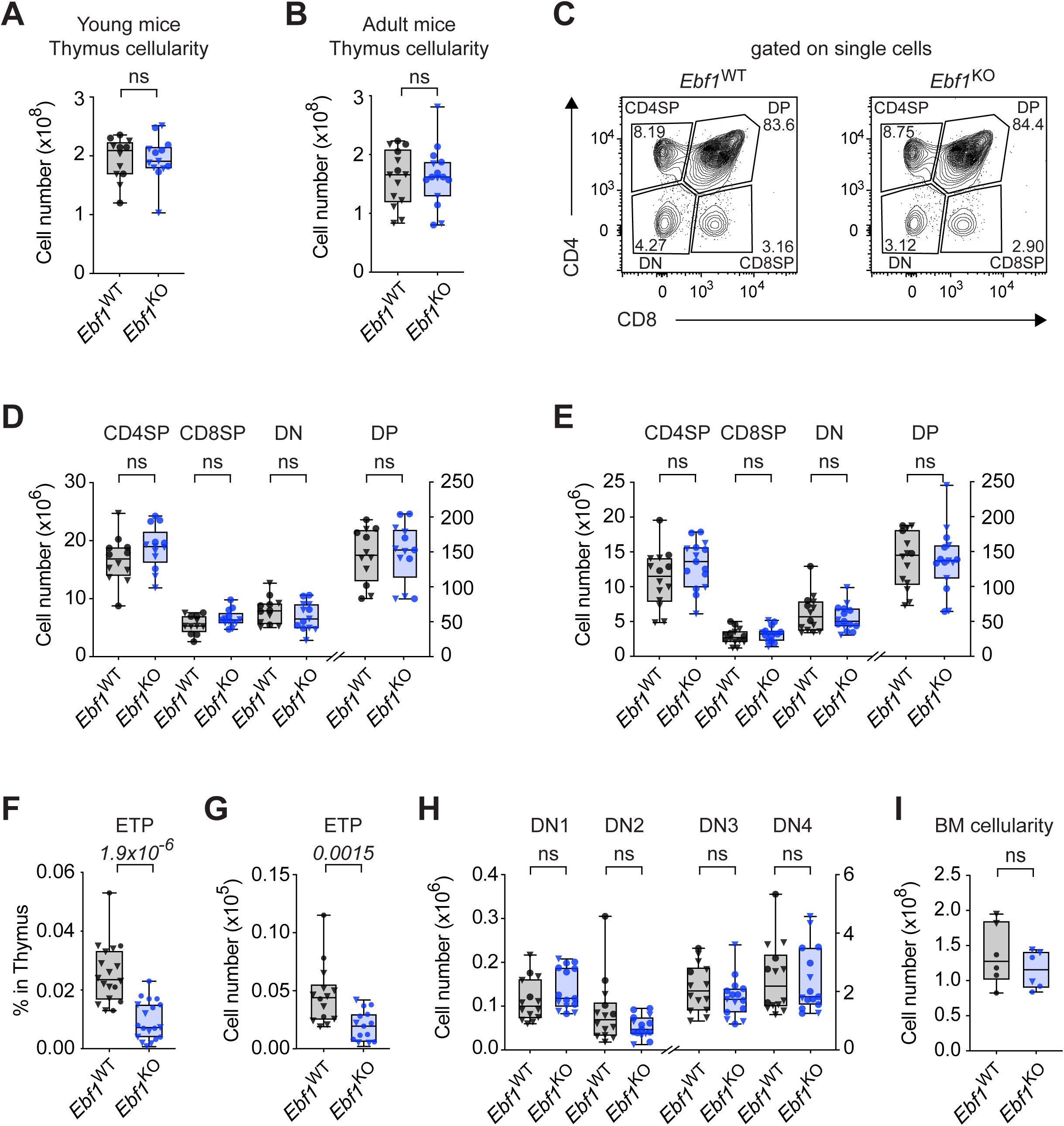
Thymocyte differentiation upon *Ebf1* deletion. Total thymus cellularity of **(A)** young mice and **(B)** adult mice. **(C)** Representative contour plots for thymocyte differentiation intermediates in *Ebf1*^WT^ and *Ebf1*^KO^ mice. Absolute number of CD4^+^ SP, CD8^+^ SP, DN and DP cells in the thymus of **(D)** young mice, and **(E)** adult mice. **(A, D)** *Ebf1*^WT^ n=12, *Ebf1*^KO^ n=13. **(F)** Total frequency and **(G)** absolute number of ETPs in the thymus of adult mice. **(H)** Absolute number of DN1-4 populations in the thymus of adult mice. **(B, E-H)** *Ebf1*^WT^ n=14, *Ebf1*^KO^ n=15. **(I)** Total bone marrow (BM) cellularity of young mice. *Ebf1*^WT^, *Ebf1*^KO^ n=6. Young mice were analyzed at 4-6-weeks-old, adult mice were analyzed at 12-weeks-old (•) represent female mice, (▼) represent male mice. Statistical significance was determined by unpaired t-test. Data are from ≥ 2 independent experiments. SP, single positive (CD4^+^ or CD8^+^); DN, double negative (CD4^-^CD8^-^Lin^-^cKit^-^); DP, double positive (CD4^+^CD8^+^).

